# Determinants of genetic diversity in sticklebacks

**DOI:** 10.1101/2023.03.17.533073

**Authors:** Mikko Kivikoski, Xueyun Feng, Ari Löytynoja, Paolo Momigliano, Juha Merilä

## Abstract

Understanding what determines species and population differences in levels of genetic diversity has important implications for our understanding of evolution, as well as for the conservation and management of wild populations. Previous comparative studies have emphasized the roles of linked selection, life-history trait variation and genomic properties, rather than pure demography, as important determinants of genetic diversity. However, these findings are based on coarse estimates across a range of highly diverged taxa, and it is unclear how well they represent the processes within individual species. We assessed genome-wide genetic diversity (*π*) in 45 nine-spined stickleback (*Pungitius pungitius*) populations and found that *π* varied 15-fold among populations (*π*_min_≈0.00015, *π*_max_≈0.0023) whereas estimates of recent effective population sizes varied 122-fold. Analysis of inbreeding coefficients (F_ROH_) estimated from runs of homozygosity revealed strong negative association between *π* and F_ROH_. Genetic diversity was also negatively correlated with mean body size and longevity, but these associations were not statistically significant after controlling for demographic effects (F_ROH_). The results give strong support for the view that populations’ demographic features, rather than life history differences, are the chief determinants of genetic diversity in the wild.

## Introduction

Understanding what determines differences in levels of genetic diversity among populations and species has important implications for our understanding of evolution, as well as for conservation and management of wildlife (DeWoody et al. 2021). According to the neutral theory of molecular evolution, genetic diversity should be proportional to population size (Kimura and Crow 1964). However, genetic diversity among populations and species varies much less than their census size, a discrepancy that has become known as the “Lewontin’s paradox” (LP; Lewontin 1974). This observation has sparked research on the determinants of genetic diversity in the wild, but the definitive answer has remained elusive (Leffler et al. 2012, Ellegren and Galtier 2016; Charlesworth & Jensen 2022).

In an influential paper, Romiguier et al. (2014) reported correlations between life history traits and nucleotide diversity across 76 animal taxa. They showed that long-lived and low fecundity species (*K*-strategists) had lower nucleotide diversities than those with short life-span and high fecundity (*r*-strategists). Their conclusions of the determinants were two-fold: on one hand, *r*-strategists have larger census population sizes, which should lead to larger effective population sizes and thereby to higher nucleotide diversity. On the other hand, *K*-strategists tend to live in more stable environments and invest more in parental care which makes them more resilient to perturbations and able to sustain populations of small size even if this entails lower genetic diversity.

Mackintosh et al. (2019) pointed out that if the conclusions of Romiguier et al. (2014) hold, and robustness to fluctuations in population size determine the nucleotide diversity, then other traits related to the ability to escape extinction, such as generalism, should correlate with genetic diversity when life history traits are overall very similar. They investigated this with 38 species of European butterflies that are all *r-*strategists. Unlike Romiguier et al. (2014) they did not find a statistically significant association between propagule size and nucleotide diversity, although they found negative correlation with body size and nucleotide diversity, or evidence for generalists having higher long-term effective population sizes. However, they discovered a positive correlation between nucleotide diversity and the number of chromosomes (a proxy of recombination rate), which was inferred as evidence for linked selection decreasing genetic diversity (Mackintosh et al. 2019). The role of linked selection limiting genetic diversity was also proposed by Corbett-Detig et al. (2015) but subsequent studies have concluded that although it influences the levels of genetic diversity (Charlesworth and Jensen 2021), linked selection may not be sufficient to explain LP (Coop 2016, Buffalo 2021).

Previous studies on determinants of genetic diversity have focused on the nucleotide diversity of species’ global metapopulations, while processes influencing nucleotide diversity of discrete contemporary populations have gained less attention (but see: Stoffel et al. 2018; Filatov 2019, Kartje et al. 2020). This is problematic, as genetic diversity across a metapopulation may simply reflect the accumulation of diversity across lineages and can be substantially higher than the diversity expected in a panmictic population with the same number of individuals (Charlesworth and Jensen, 2022). By studying nucleotide diversity in 45 populations of the nine-spined stickleback (*Pungitius pungitius*), we aimed to address this knowledge gap.

The nine-spined stickleback is a small teleost fish with circumpolar distribution, occupying both marine and freshwater habitats. Previous studies have demonstrated life history differences between marine and freshwater environments; individuals in freshwater habitats, especially in ponds, are bigger, mature later and have longer life-span than those in marine environments (Herczeg et al. 2009, DeFaveri et al. 2014). Freshwater individuals also have bigger clutches, bigger eggs and hatch larger offspring (Herczeg et al. 2010). However, on the scale of the whole animal kingdom, the life history differences are subtle. In addition to those characteristics, populations differ in the sizes of their living area (*viz*. sea, lake, pond), connectivity (*viz*. sea, connected streams and lakes, landlocked ponds) and their evolutionary past, namely most of the freshwater populations have been established and undergone local adaptation to their environment after the last glaciation.

With access to high quality whole-genome re-sequencing data from a large number of replicate populations, nine-spined sticklebacks provide a suitable model system for exploring within species variation in nucleotide diversity and its determinants. We hypothesized that if life history traits are the major determinants of genetic diversity, their influence on genetic diversity should exceed that of the populations’ demographic history. Conversely, would the populations’ demographic history be the major determinant of genetic diversity, one would expect to see various indicators of population demography (*viz*., level of inbreeding, population size bottlenecks and contemporary effective population size) would consistently and better explain intraspecific variation in nucleotide diversity as compared to variation in life history traits.

## Results

We whole-genome sequenced 892 individuals from 45 nine-spined stickleback populations across the northern hemisphere (Fig. 1A,B) to the depth of 10-20X, mapped the data to the reference genome (Varadharajan et al. 2019) and called the variants following the GATK4 best practices (McKenna et al. 2010; see Methods). The autosomal part of the 404 Mbp assembly contained in total 23,352,704 single nucleotide polymorphisms (SNPs). We calculated the genome-wide nucleotide diversity (*π*) estimates for the entire autosomal genome and, to allow comparisons to previous studies, for silent site substitutions in exonic regions only.

**Figure 1.**
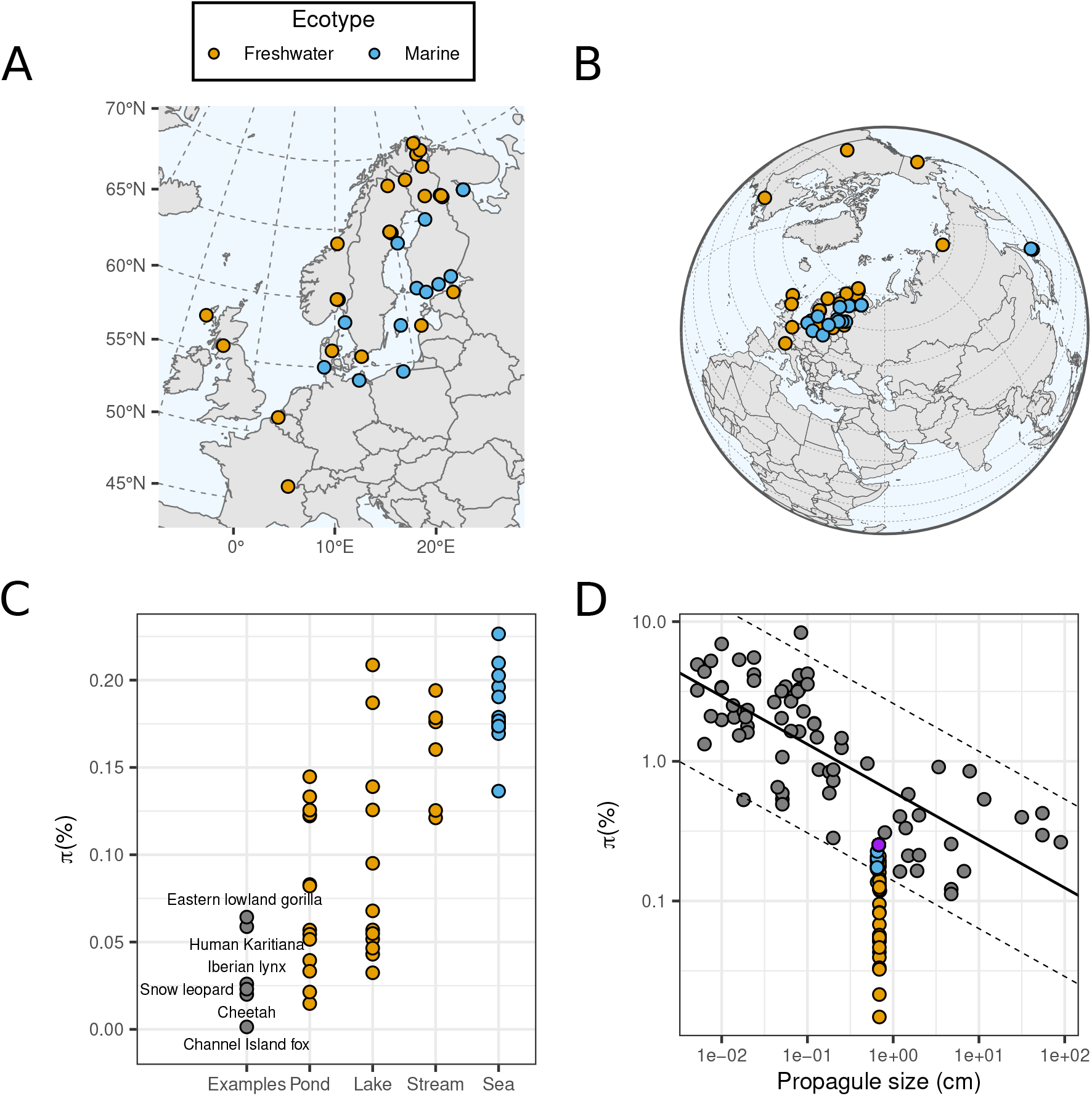
Genome-wide nucleotide diversity (*π*) in *P. pungitius* populations. (A, B) Maps of the 45 study populations. (C) Estimates of *π* in *P. pungitius* populations (colored dots) in comparison to selected species with exceptionally low genetic diversity (gray dots, Robinson et al. 2016). (D) *π* as a function of propagule size in 45 nine-spined stickleback populations (orange and blue dots) projected on the results of Romiguier et al. 2014 (gray dots). Purple dot shows the nucleotide diversity for a global set of nine-spined sticklebacks (n=892). Solid and dashed lines indicate the linear model fitted to Romiguier et al. (2014) data and the standard error of the model (±1.96-times the standard deviation of residuals), respectively.Here, the *P. pungitius* fry length (6.5–6.9 mm, Herczeg et al. 2010) is used as a proxy of propagule size.

The population-level estimates of *π* was found to vary between 0.00015 and 0.0023 (15-fold difference) being lowest in the landlocked pond population (Pyöreälampi, Finland) and highest in a Japanese marine population (Biwase Bay, Hokkaido; Fig. 1C). Overall, the marine populations had higher *π* than the freshwater populations (Welch’s t-test, t_43_=-7.24, p<1.010^−8^) and *π* varied much less among the marine than freshwater populations (Fig. 1C). The global *π* (0.0025) was higher than *π* for any of the individual populations, showing that genetic diversity of a global metapopulation can be very different from those of discrete populations (Fig. 1D). While the global *π* fitted well to the linear model Romiguier et al. (2014) reported for nucleotide diversity and propagule size, all populations were below their predictions and most were outside of the standard error range indicating poor fit to the model (Fig. 1D). Nucleotide diversity in freshwater populations was uncorrelated (*F*_1, 26_=0.47, *p=* 0.50, Fig. S1) with the habitat area, a proxy of census population size (*N*_*C*_).

Based on the *π* estimates, we calculated the long-term effective population sizes *N*_*e*_^*π*^ according to the neutral theory expectation *N*_*e*_^*π*^ = *π/*4μ (Kimura 1991), assuming mutation rate per generation 7.06 × 10^−9^(Chaowei Zhang, unpublished). However, *N*_*e*_^*π*^ alone provides very little information of past demographic events as the historical *N*_*e*_ can vary across time. To get insight to this, we estimated the historical *N*_*e*_ with MSMC2 (Schiffels & Ke 2020). Ignoring the first two time segments, typically unreliable due to the small amounts of information available, the *N*_*e*_ trajectories showed substantial variation both among populations and across time within a given population (Fig. 2A). We considered the most recent time segment provided by MSMC2 as a proxy of the (near) contemporary effective population size, *N*_*e*_^*MR*^. A comparison of long-term and contemporary *N*_*e*_ estimates showed that the *N*_*e*_^*π*^ estimates were 3–31 times higher than those for *N*_*e*_^*MR*^, while the range of *N*_*e*_^*π*^ (15-fold) was significantly smaller than the range of N_e_^MR^(122-fold). The ratio of the *N*_*e*_^*π*^ and the *N*_*e*_^*MR*^ was negatively correlated with *π* (*r* = -0.75; Fig. 2B) showing that populations with the lowest genetic diversity display the greatest differences in the *N*_*e*_^*π*^ and *N*_*e*_^*MR*^ estimates and thus drive the difference seen in the range of the values. In all 45 cases, the *N*_*e*_^*π*^ exceeded the *N*_*e*_^*MR*^ indicating that, at least in our study populations, the diversity created by a historically larger population size or past demographic events surpasses that from more recent times.

**Figure 2.**
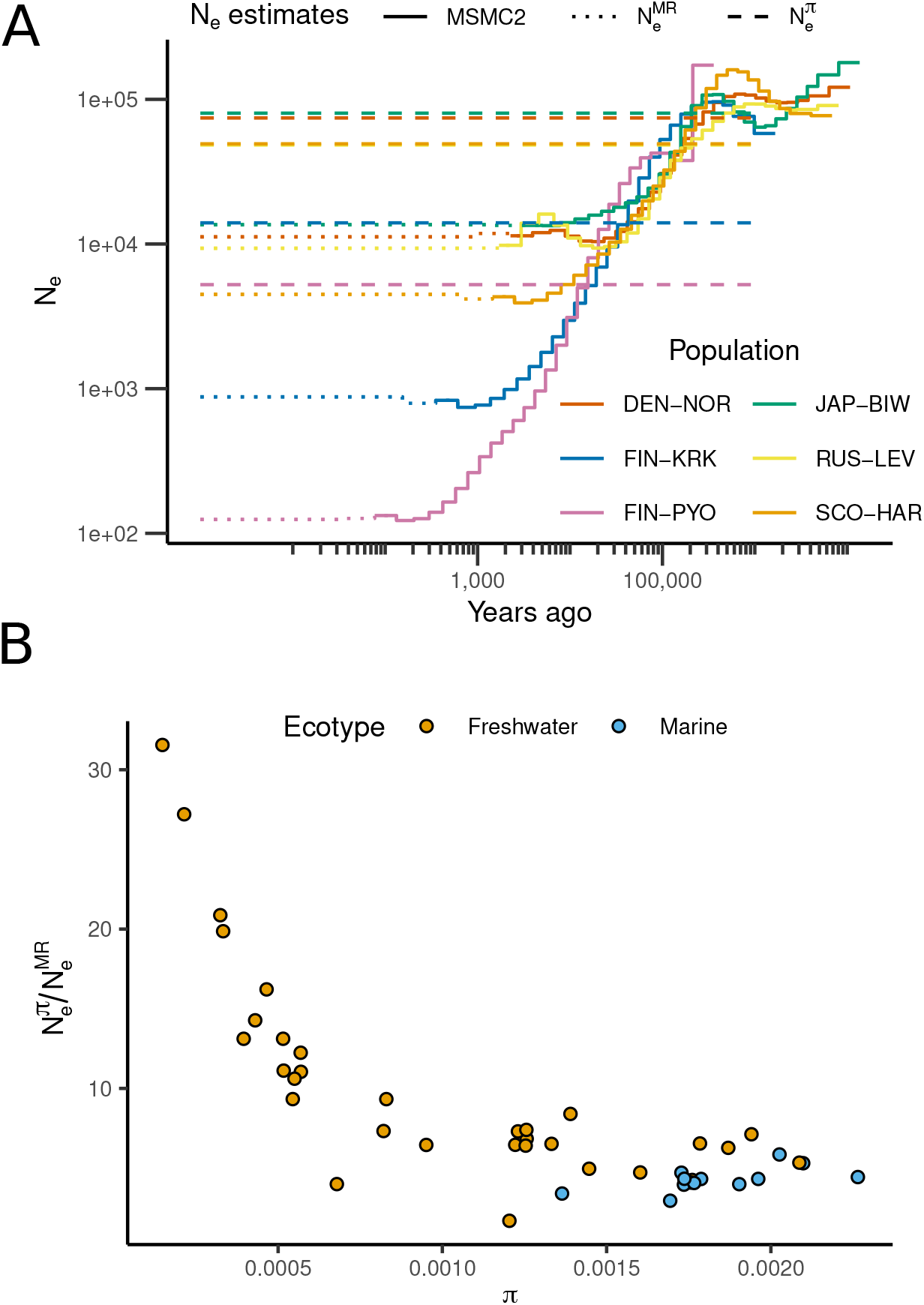
Effective population sizes of nine-spined stickleback populations across time. (A) Contemporary and historical N_e_ (dotted and solid lines, respectively) estimated with MSMC2 and the long-term effective N_e_ (dashed line) calculated from *π* (N_e_^π^= *π*/4*μ*) for six study populations representing variable demographic histories. (B) The ratio of long-term and contemporary N_e_ s (y-axis) and the genetic diversity (*π*; x-axis) across the 45 populations are negatively associated, largely due to high N_e_ ratio in populations with low *π*.

Ceballos et al. (2018) showed that the runs of homozygosity (ROH) within genomes reflect the individuals’ demographic histories. We used BCFtools roh (ver. 1.16, Danecek et al. 2021) to infer ROHs within every individual and projected the resulting number of homozygous regions (N_ROH_) on their total length (S_ROH_). The distribution of the estimates revealed a great deal of variability in demographic histories and allowed dividing the individuals and populations into six groups with k-means clustering (Fig. 3A; Ceballos et al. 2018). While the marine populations were mainly close to the origin indicating admixture and large effective population sizes, the freshwater populations showed highly variable histories with varying levels of bottlenecks and inbreeding (Fig. 3A). More specifically, the patterns of ROHs allowed separating large populations from medium-sized populations, and consanguineous populations from bottlenecked populations, leaving both bottlenecked and consanguineous populations from small ponds as extreme outliers (Fig. 3A). The effect of the demographic history on nucleotide diversity became evident when we related the *π* to the levels of inbreeding (F_ROH_): across all individuals, nucleotide diversity displayed a strong negative association with the genomic inbreeding coefficient (Fig. 3B).

**Figure 3.**
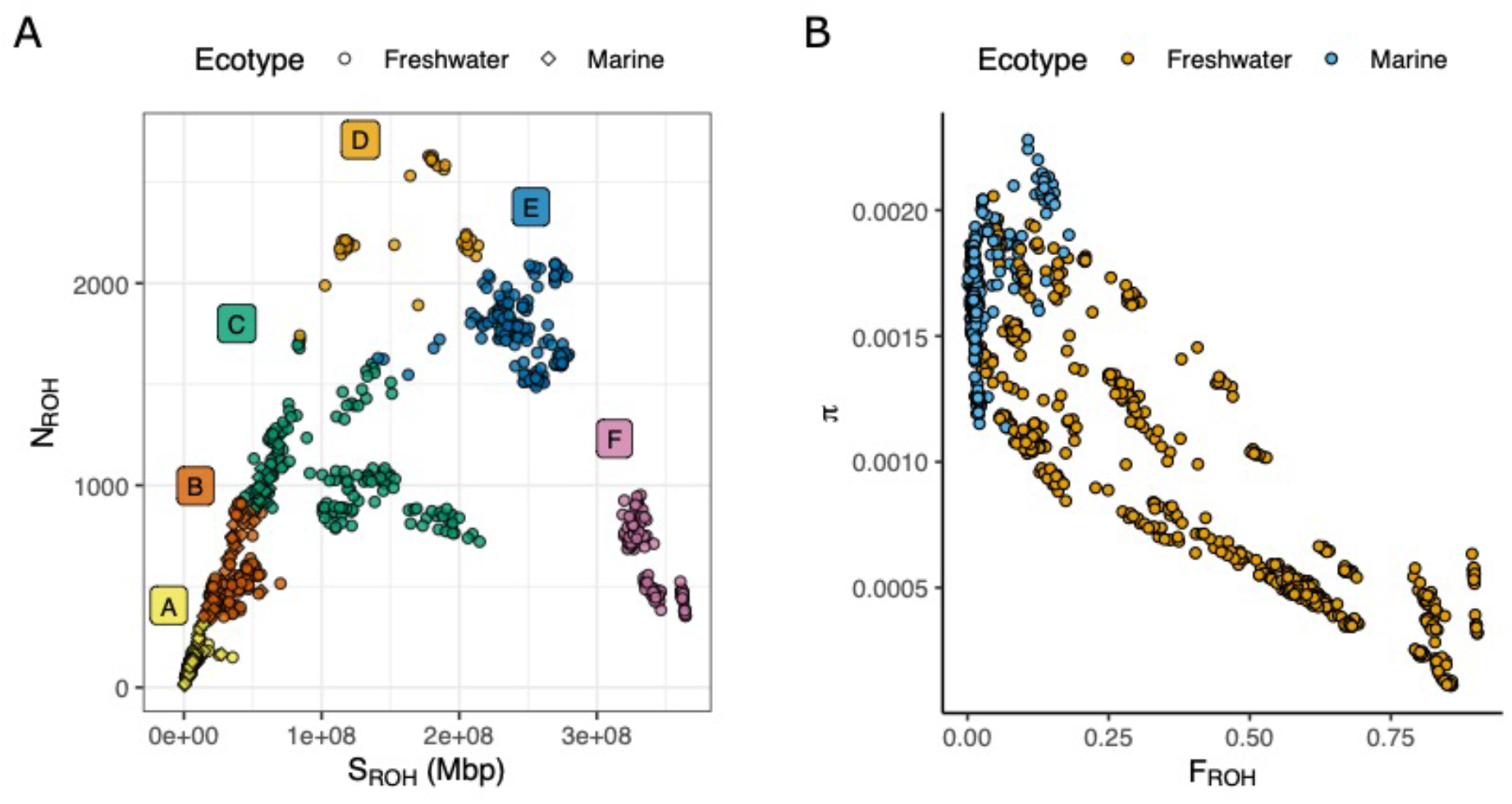
Analysis of runs of homozygosity in individual nine-spined sticklebacks. (A) Association between the number of homozygous stretches in the genome (N_ROH_) and their total length (S_ROH_). Individuals from populations with similar demographic histories are clustered and can be divided into six categories (A–F) following Ceballos et al. (2018): A=admixed populations, B=large populations, C=medium-sized populations, D=consanguineous populations, E=bottlenecked populations, F=bottlenecked and consanguineous populations. (B) The relationship between nucleotide diversity and homozygosity (F_ROH_).

To assess effects of life-history traits on *π*, we used data on longevity and adult body size available for part of the populations (5 marine and 8 freshwater populations). To study this, we fitted linear mixed–models with *π* as the response variable, longevity, adult standard length, and F_ROH_ as fixed effects. Phylogenetic non-independence was taken into account by incorporating a phylogenetic correlation matrix as a random effect into the model (see Methods). We specified two models, one with the ecotype (marine/freshwater) as a fixed effect without interaction terms with ecotype and other fixed effects. The other model included the interactions of ecotype and other fixed effects. The models were fitted with MCMCglmm R package (ver. 2.34 Hadfield 2010). For both models, two different priors were used (see Methods), yielding altogether four models (two priors, two models). The best model was selected based on the lowest deviance information criterion (DIC).

The model with lowest DIC was the one without the interactions of ecotype and other fixed effects. *π* was not statistically significantly associated with longevity and body size, whereas the effect of F_ROH_ on *π* was negative and statistically significant (Table 1).

**Table 1.** Results of Bayesian linear model fitted with MCMC algorithm (Markov Chain Monte Carlo) with population’s *π* as a response variable, and habitat, F_ROH_, bodysize, and longevity as explanatory variables.

| Parameter | Posterior mode | $p_{MCMC}$ | 95% HPDI |
| --- | --- | --- | --- |
| Intercept (FW) | -0.509 | 0.33 | -1.03 – 0.60 |
| Intercept (M) | 0.249 | 0.74 | -0.74 – 0.93 |
| $F_{ROH}$ | -0.684 | 0.002 | -1.09 – -0.34 |
| Body size | -0.019 | 0.79 | -0.44 – 0.35 |
| Longevity | 0.050 | 0.86 | -0.35 – 0.38 |

FW=Freshwater, M=Marine
$p_{MCMC}$ =two-times the posterior probability mass above zero, small p-values indicating statistically significant difference to zero.
HPDI=Highest posterior density intervals. Interval not spanning zero indicates significant difference to zero at 5% confidence level

## Discussion

Lewontin and Hubby’s (1966) pioneering work on protein gel electrophoresis revealed unexpectedly large amounts of genetic variation in natural populations and led to a paradigm shift in the theory of molecular evolution (Kimura 1977). Despite decades of research with increasingly sophisticated measurement and analysis methods, the original finding of their study, surprisingly similar levels of diversity across species with highly unequal population sizes – known as Lewontin’s paradox (LP) – remains a puzzle. While the early attempts of resolving LP emphasized demographic explanations (e.g., population bottlenecks; Kimura 1984; Ohta & Kimura 1973; Nei & Graur 1984), much of the recent work has focused on linked selection (e.g. Corbett-Detig et al. 2015; Coop 2016; Buffalo 2021; Charlesworth & Jensen 2021) before returning to demographic factors, especially recent population size expansions (Charlesworth & Jensen 2022) implicitly recognized also in studies of range expansions (Ramachandran et al. 2005). Several studies applying modern high-throughput DNA sequencing have correlated levels of nucleotide diversity with biological differences among species (Romiguier et al. 2014, Corbett-Detig 2015, Mackintosh et al. 2019). However, genome-wide studies focusing on differences among populations of single species have remained limited (but see Filatov 2019, Kartje et al. 2020).

Our analyses of 45 populations of nine-spined sticklebacks show that genome-wide genetic diversity (*π*) varies by an order of magnitude (15-fold) among the populations and that demography, rather than life history differences, appear to be the chief determinant of contemporary diversity. The observation of higher *π* in marine than in freshwater populations conforms to the general pattern in fishes (Gyllensten 1985, Ward et al. 1994, DeWoody and Avise 2000, Ward 2004, McCusker and Benzen 2010, Merilä 2014, but see Manel et al. 2020) and can be understood in the light of markedly different demographic histories and N_e_ s of marine and freshwater populations (Fig. 2 & 3) as well as prevalence of recent admixture in the marine environment (Yamasaki et al. 2020, Feng et al. 2022). While the *π* of marine populations was generally high, the demographic footprints among the different freshwater populations were highly diverse, ranging from marine-like demography in riverine and larger lake populations to extreme signs of population bottlenecks and inbreeding in isolated ponds.

Although we found negative associations between two life history traits (body size and longevity, Fig. S2) and *π*, these effects were not statistically significant when the runs-of-homozygosity (ROH) derived inbreeding coefficient (F_ROH_), a proxy of demographic effects, was added as a covariate. In contrast, the strong negative correlation between *π* and F_ROH_ across the populations gives further support for the differences in *π* being driven by demography. Our study populations reproduced the patterns predicted by Ceballos et al’s (2018) for N_ROH_ *vs*. F_ROH_ under different demographic scenarios with one exception: our data additionally included populations that were both extremely bottlenecked and inbred and as a result, contained relatively few but extremely long homozygous genome stretches (“Group F” in Fig. 3A). The astonishingly low SNP densities (≈100 SNPs/1 Mb) in some pond populations, originated after the last glacial maximum ca. 10,000 years ago and probably isolated ever since, are among the lowest observed in vertebrate species, coming close to that of the island foxes in Baja California regarded as the most genetically depauperate vertebrate known to science (Robinson et al. 2016).

Negative correlations between body size and longevity and *π* have been found across a wide range of animal taxa (Romiguier et al. 2014, Brüniche-Olsen et al. 2018, Mackintosh et al. 2019) and are taken as support for the body size limiting population density (White et al. 2007) and thereby long-term N_e_. While it is conceivable that the size and longevity of nine-spined stickleback populations could influence their demography and thereby *π* (cf. Romiguer et al. 2014), correlation does not always imply causality; we find it more likely that low *π* in the smallest ponds occupied by large and long-lived fish is caused by strong population size bottlenecks and inbreeding.

We computed a proxy of the contemporary effective population size *N*_*e*_^*MR*^ using the program MSMC2 and the long-term effective population size *N*_*e*_^*π*^ from the genome-wide *π*. The discrepancy between the two estimates suggests that the populations are not in drift–mutation equilibrium and contain old segregating variation that differently affects the two approaches. The difference between the estimates *N*_*e*_^*MR*^ and *N*_*e*_^*π*^ was greatest (31-fold) in the inbred pond population with the lowest nucleotide diversity, while the estimates for marine populations were more similar (3-6 fold difference). This is interesting in the view that theory predicts (e.g. Galtier & Rouselle 2020) that, following a reduction in population size, populations should approach equilibrium in a number of generations on the order of the post-reduction *N*_*e*_. Hence, pond populations should approach equilibrium values much faster than the marine populations with large *N*_*e*_, yet the *π* estimates of the pond populations reside farther away from the equilibrium values inferred with MSMC2. One possible explanation for this is that, even if demography generally is the strongest determinant of genetic diversity, in extremely small populations other factors–such balancing selection and selection against lethal recessive alleles become significant.

Our findings challenge some of the conclusions by Romiguier et al. (2014). They observed that the range of genetic diversities (0.1% to 8.3%) across animal taxa had a strong taxonomic effect and closely related species tended to have similar genetic diversities. They reasoned that ‘such a strong taxonomic effect would be unexpected if stochastic disturbances and contingent effects were the main drivers of genetic diversity, because species from a given family are not particularly expected to share a common demographic history.’ While this sounds logical, their study design did not allow evaluating how demography below species level contributed to genetic diversity and was thus biased towards finding evidence for species biology and ecology as determinants of genetic diversity. As shown by our analyses (Fig. 1D), the metapopulation level estimates effectively hide much of the demographically driven variance in nucleotide diversity and can be substantially larger than they would be in a panmictic population of the same size (Löytynoja et al. 2023).

In conclusion, analyses of extensive whole-genome data from 45 stickleback populations point to the importance of demography as a determinant of genetic diversity across populations. As such, our observation of the range of recent effective population sizes being eight times larger than that of genome-wide diversities (122-fold vs. 15-fold) is consistent with the conclusions of the recent review by Charlesworth & Jensen (2022) emphasizing the role of past demographic events as a major determinant of genetic diversity. The 15-fold range of genetic diversities found within a single species is astonishing, drawing attention to the extremely low genetic diversities in bottlenecked and consanguineous pond populations. The discrepancy between *N*_*e*_^*MR*^ and *N*_*e*_^*π*^ was the greatest in these tiny populations, and given that their isolation probably greatly exceeds 4*N*_*e*_ generations (Hartl and Clark 2007), their elevated levels of diversity must be retained by factors other than demography. However, it is clear that the excess diversity of the small pond populations originated during the historical large population size, emphasizing that the mechanisms creating genetic diversity and those retaining it under altered circumstances can be different.

## Acknowledgements

Kirsi Kähkönen and Miinastiina Issakainen are thanked for their help with laboratory work and numeous collaborators and colleagues for their help in obtaining the samples (listed in Acknowlegedments of Feng et al. 2022). This research was supported by LUOVA doctoral programme (funding to MK), Academy of Finland (#129662, 134728 and 218343 to JM, 322681 to AL), grant from Helsinki Institute for Life Sciences (HiLife; to JM) and Alfred Kordelin Foundation (grant #210190 to MK). The work described in this paper was also partially supported by a grant from the NSFC/RGC Joint Research Scheme sponsored by the Research Grants Council of the Hong Kong Special Administrative Region, China and the National Natural Science Foundation of China (Project No. N_HKU763/21). CSC – IT Center for Science, Finland, is acknowledged for access to computational resources.

## Methods

### Data acquisition

The study utilized 892 whole-genome re-sequenced samples from 45 locations covering much of the species distribution range in Europe, Siberia, Japan and North America (see Supplementary Table S1 for sampling sites and their coordinates). The sample collection and associated metadata have been described previously in Feng et al. (2022). In brief, samples were collected between 2006 and 2015 using seine nets and minnow traps during the local breeding seasons in accordance with legislation in the given locality. Ethanol preserved specimens or fin-clips were brought to the laboratory in Helsinki for DNA extractions. DNA was extracted with the standard phenol-chloroform (Sambrook and Russell 2006) and salting out methods (Miller et al. 1988, Bruford et al. 1998). The quality and quantity of the extracted DNA were assessed with NanoDropTM 2000 spectrophotometer (Thermo Fisher Scientific), Qubit fluorometer (Qubit® 2.0 Invitrogen, Qubit™4 Thermo Fisher Scientific) and gel electrophoresis.

### Whole-genome re-sequencing and variant calling

Whole-genome DNA sequencing was performed with Illumina short-read protocols (HiSeq 2500/4000 instruments) and carried out in Beijing Genomics Institute (BGI, Hong Kong SAR, China), Novogene (UK) Company Limited, and the DNA Sequencing and Genomics Laboratory, University of Helsinki (Helsinki, Finland). Sequencing depth of the samples varied between 10X and 20X (Feng et al. 2022).

Whole-genome re-sequencing reads were aligned to the nine-spined stickleback reference genome ver. 6b (Feng et al. 2022), a modified version of the ver. 6 genome (Varadharajan et al. 2019) with BWA-mem algorithm (Li 2013). Alignments were further processed by marking the duplicate reads with SAMtools (Li et al. 2009) and performing local realignment with the Genome Analysis Toolkit (GATK, ver. 4.0.1.2, McKenna et al. 2010). The variants were called jointly for the whole data set with the GenotypeGVCFs command in GATK. Only biallelic SNPs were retained.

Four samples (CAN-TEM-16, CAN-TEM-18, SWE-LUN-46, and SWE-LUN-39) had high rates of missing data and they deviated from the other samples from the same populations in runs-of-homozygosity analyses and nucleotide diversity. Therefore, they were excluded from the further analyses.

### Estimation of nucleotide diversity (π)

Nucleotide diversity estimates were obtained in two ways. First, the autosomal nucleotide diversity (*π*) was estimated from the called genotypes with the vcftools (ver. 0.1.16, Danecek et al. 2011) command --site-pi, by dividing the total with the length of the genome excluding the linkage group 12 (sex chromosomal linkage group in *P. pungitius;* Rastas et al. 2016). *π* was calculated both at the population and individual level. In order to assess the *π* calculated from the called variants, *π* were also estimated from site frequency spectra (SFS; see Momigliano et al. 2021). Folded SFSs were estimated from read alignments (the bam files) using ANGSD (ver. 0.921-3-g40ac3d6, Korneliussen et al. 2014). A positive mask (Kivikoski et al. 2021) was first applied to prune the sites based on genome mappability. For each population the following filters were applied: biallelic sites with no missing data were retained, sites with mapping quality and base quality below 20 were removed (-minMapQ and -minQ option in ANGSD), and further requirement was that each remaining site has no heterozygote counts more than 70% of all counts. The resulting *π* had a correlation of 0.998 with those estimated from the variant calls.

Key previous studies in determinants of genetic diversity (e.g. Romiguier et al. 2014) have been based on RNA-sequencing data, whereas the present study utilizes whole-genome re-sequencing. In order to derive *π* estimates comparable to such studies, we aimed to replicate the calculation of *π* replicating the analysis of Romiguier et al. (2014). First, the variant calls were converted to fasta sequences using the vcf2fasta.py script from Rousselle et al. (2020). This step was performed for single copy BUSCO genes that were identified as follows. The gene annotation (Varadharajan et al. 2019) was cleaned by merging the autosomal gene annotations following the separation of the overlapping and non-overlapping genes. For overlapping gene annotations, we then searched from the overlapped genes to see if any of them can completely cover the merged entry. If this type of long entry existed, then this entry/gene annotation was kept. This step created the single copy gene list. The single copy gene list was further compared with the complete single copy BUSCO genelist (Varadharajan et al. 2019). If any BUSCO entry overlapped with more than one single copy gene, then this BUSCO entry was discarded. Only BUSCO entries uniquely overlapping with single copy genes were kept. This step resulted in 3209 complete single copy BUSCO genes. The synonymous and non-synonymous changes and the *π* based on those were calculated with the dNdSpiNpiS_1.0 script (URL: https://kimura.univ-montp2.fr/PopPhyl/index.php?section=tools; see Romiguier et al. 2014).

Observed nucleotide diversities were compared to those reported from other populations and organisms (see Fig. 1C). The *π* for cheetah, snow leopard, human and eastern lowland gorilla were obtained from the supplementary data S1 of Robinson et al. (2016). Original values are reported in Dobrynin et al. (2015), Cho et al. (2013), Meyer et al. (2012), and Xue et al. (2015), respectively. For the Channel Island fox, the *π* was obtained from Robinson et al. (2016) and for the Iberian lynx form (Abscal et al. 2016)

### Analysis of runs of homozygosity and admixture

Runs of homozygosity were analyzed with the BCFtools programme’s roh command (ver. 1.16, Narasimhan 2016). The command was run with the following parameters: *-G30 -I*, and providing the nine-spined stickleback linkage map (*-m* flag, Kivikoski et al. 2023) to utilize correct recombination rates. ROH segments longer than 20 kbp were retained. Individuals were divided in six demographic history groupings as in Ceballos et al. (2018). The grouping was performed with k-means clustering (*kmeans* function on R version 4.2.1, setting the number of clusters to six) based on individuals’ total length of homozygous segments (S_ROH_) and mean number of homozygous segments (N_ROH_).

### Estimating the effective population size and census size

The contemporary and long term effective population sizes (*N*_*e*_) were estimated with MSMC2 (Malaspinas et al. 2016) following Schiffels and Wang (2020) using default settings. The mutation rate per generation is 8.94 × 10^−9^ for pond populations and 5.17× 10^−9^ for marine populations (Chaowei Zhang, unpublished), and here the average 7.06 × 10^−9^ was used for all populations. Based on DeFaveri et al. (2014), the generation length was defined as 2.5 and 3.2 years for marine and freshwater populations, respectively. The surface area of the habitat was used as a proxy of the census size (*N*_*C*_). For the freshwater populations, the populations were assumed to occupy the entire habitat.

### Life-history traits

Data on body size and longevity were available for 13 study populations from DeFaveri et al. (2014, see DeFaveri et al. 2015 for the data). For these populations, the mean standard length of the five largest males and females per population were used as a proxy of body size. Length of the fish can vary among the sample collections (Arai et al. 2010) for example due to variation in age structure (Raeymaekers et al. 2008, DeFaveri et al. 2014) and this approach reduces potential bias due to differences in population age structure (Forsman 1991). The maximum age (years) per population was used as a proxy of longevity. The compiled life history data used in this study is listed in Supplementary table S1.

### Linear models assessing the effects of life history traits and demographic histories on π

Effects of life history traits (adult standard length and longevity) and demographic history (homozygosity, F_ROH_) on *π* were assessed with linear mixed-models accounting for the phylogenetic non-independence of the populations and individuals (see e.g. Buffalo 2021). Eight freshwater populations and five marine populations with available life history information were used. The models were fitted with the MCMCglmm package (ver. 2.34, Hadfield 2010), providing the phylogenetic correlation matrix (see below) as a random effect.

We analyzed the data at the population level, and two kinds of models were fitted: a model with interaction between the ecotype (freshwater/marine) and the fixed effects (model 1), and a model with ecotype as a fixed effect, but without the interactions (model 2). In both models, population’s *π* was the response variable and the explanatory variables were the longevity, adult standard length and the arithmetic mean of the F_ROH_ values obtained from the individuals of the population. All variables were mean-standardized before the model fitting. We fitted the models with two uninformative priors, such that:

prior A: list(R=list(V=1, nu=1), G=list(G1=list(V=1, nu=1, alpha.mu = 0, alpha.V = 1000)))

prior B: list(R=list(V=1, nu=0.002), G=list(G1=list(V=1, nu=0.002)))

From the four fitted models (two models, two priors per model), the model with the lowest deviance information criterion (DIC) was selected (model 2 with prior B).

The phylogenetic correlation matrix was derived by calculating the identity-by-state distance of the individuals from the LG4 (chromosome 4, longest autosomal chromosome) VCF files using plink (ver. 1.90, Purcell et al. 2007), calculating the rooted (*Pungitius tymensis* as an outgroup) neighbor-joining tree with R package ape (ver. 5.6, Paradis and Schliep 2019) and converting that to a covariance matrix with vcv and cov2cor functions, respectively. For the population level model, the matrix was subsetted so that it included one randomly sampled individual per population.

## Data availability statement

The whole-genome re-sequencing data have been published previously by Feng et al. (2022) and they are available in the European Nucleotide Archive (ENA) (https://www.ebi.ac.uk/ena) under accession code PRJEB39599 and Zenodo Open Repository: https://zenodo.org/record/6951309, respectively. The life-history data were compiled from an earlier study by DeFaveri et al. (2014) and provided in supplementary Table S1, phylogenetic correlation matrix is provided as a supplementary file. Linkage map used in the ROH analyses were obtained from Kivikoski et al. (2023; https://github.com/mikkokivikoski/InverseMappingFunctions/tree/main/Data).

