## Supplemental figures for "Determinants of genetic diversity in sticklebacks"

<u>Content</u>

Supplementary table S1: Tab-separated table (separate file) containing the information of the study populations (sample size, locality), life history (when available), estimated nucleotide diversity ($\pi$), ROH, and effective population size estimates.

Supplementary table S2: Phylogenetic correlation matrix used in the linear-mixed models (separate file).

Supplementary figures S1-S2

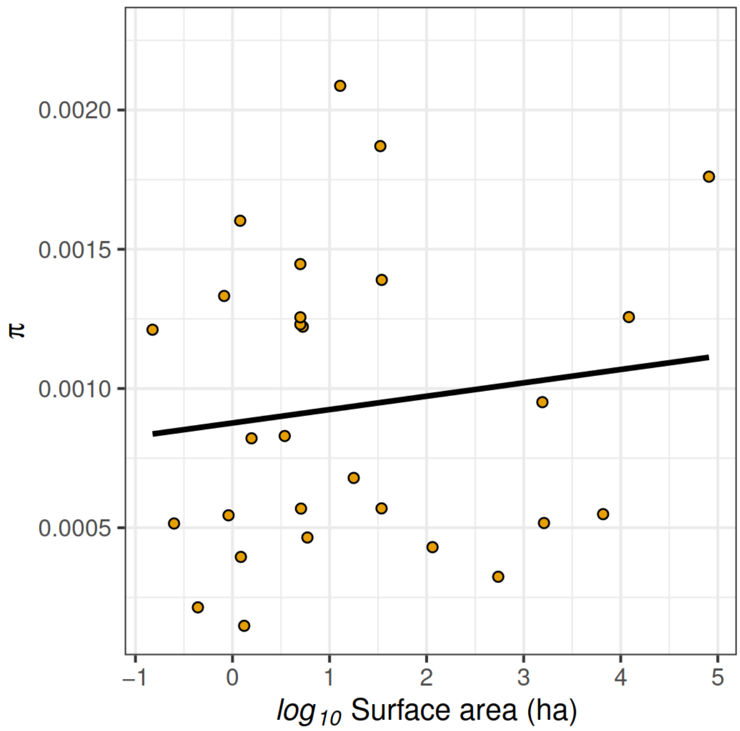


**Supplementary Figure S1.** *π* as a function of surface area in pond and lake populations. The positive association was not statistically significant ($F_{1, 26}$=0.47, *p*= 0.50)

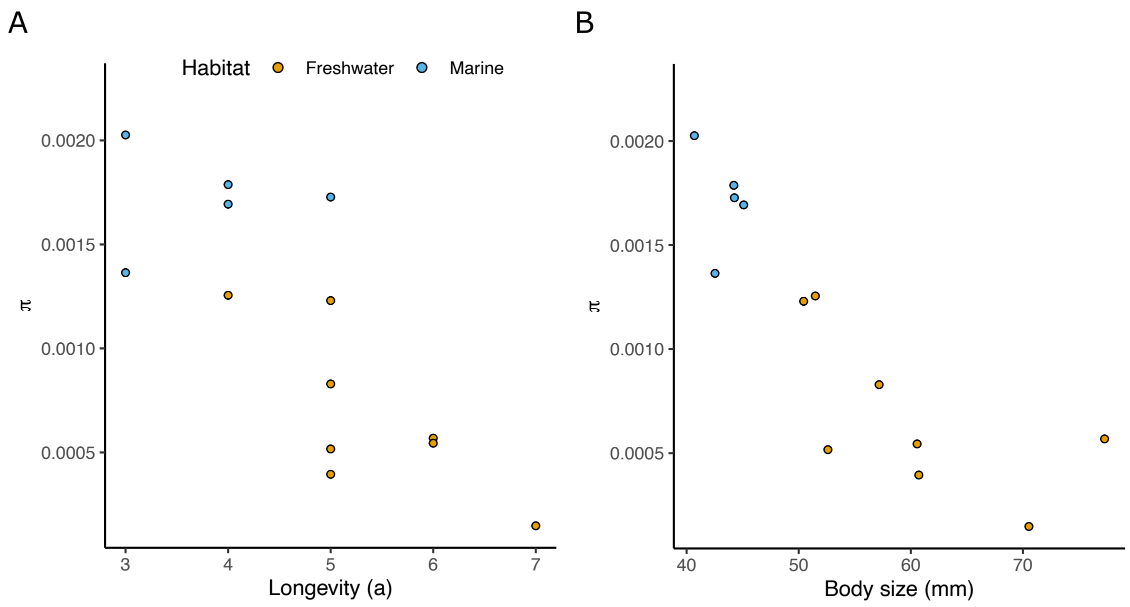


**Supplementary Figure S2.** Population *π* as a function of life-history characters, longevity (A) and body size (B).
